# Making the best of a sticky situation: Infection-mediated endothelial activation promotes new interactions with adhesins of the host-adapted Lyme disease spirochete

**DOI:** 10.64898/2026.02.20.707099

**Authors:** Daiana Alvarez Olmedo, Xi Tan, Colton Scott, Aleksandra Shcherbakova, George Chaconas

## Abstract

Lyme disease, caused by the spirochete *Borrelia burgdorferi* and closely related Lyme *Borrelia*, is the most prevalent tick-borne illness in the northern hemisphere. An important pathway for *B. burgdorferi* dissemination is its interaction with, and traversal of the vascular endothelium, a process that is not well understood and is mediated by spirochete surface adhesins. We show here that infection-induced activation of the endothelium in BALB/c mice results in new *B. burgdorferi*-endothelial interactions, indicating the presence of spirochete factors that interact specifically with activated endothelial cells. We show that these interactions are mediated by spirochetal surface proteins whose synthesis is dependent upon *B. burgdorferi* host adaptation.

We used intravital microscopy and a functional gain approach to assess the binding of spirochetes that withstand the shear force of blood flow in post-capillary venules of living mice. We identified five previously undescribed, shear force–resistant adhesins that selectively mediate binding to activated endothelium (BBA66, P66, BBA36, BBA07 and DbpA) and interact with activation-induced endothelial surface changes. Two of these adhesins (P66 and DbpA) have been implicated in the spirochete extravasation process. We also identified seven previously undescribed shear force-resistant adhesins that target pre-activated endothelium (BBA04, BBK53, BBK07, BBA65, BB0844, ErpK and OspC). Three of these (BBK53, ErpK and OspC), display reduced binding to activated endothelium, a property that may facilitate the multi-step pathway of vascular transmigration. In particular, OspC has been previously implicated in spirochete extravasation.

In summary, our results reveal a dynamic interaction network between the spirochete and the endothelium where the spirochete capitalizes on activation of the endothelium to establish new interactions and at the same time disrupt others. We propose that this scenario is part of a sequential interaction network leading to transendothelial migration of the spirochetes and subsequent tissue invasion. This work opens a new area of study focusing on eleven new adhesins described here and their vascular interactions and role in spirochete extravasation.

## Introduction

Lyme disease, a multi-organ zoonosis with an expanding range is commonly encountered in the northern hemisphere (Steere et al., 2016; Stanek and Strle, 2018; Coburn et al., 2021; Radolf et al., 2021; Mead, 2022; Smith, 2025). It is caused by the bacterial pathogen *B. burgdorferi*, and related species. In humans and other vertebrates, the infection process is mediated by infected hard-shelled *Ixodes* ticks that inoculate the spirochetes while taking a blood meal. The disease can result in serious consequences if not diagnosed and treated quickly.

The evolutionary interactions between pathogens and hosts have spawned escalating strategies to both promote and combat infections. This has resulted in intricate networks of offensive and defensive strategies. It has long been known that *B. burgdorferi* can promote endothelial activation in endothelial cells grown in culture (Sellati et al., 1995; Sellati et al., 1996; Wooten et al., 1996; Burns et al., 1997; Burns and Furie, 1998; Lochhead et al., 2012). Recently, in our intravital imaging experiments, we noted that infection with the Lyme disease spirochete results in endothelial activation through cytokine signalling in the mouse (Tan et al., 2021). Host neutrophils appear to be the predominant source of the cytokines. Cytokine release is promoted by the *B. burgdorferi* surface proteins P66 and DbpAB (Tan et al., 2023) resulting in the upregulation of a variety of endothelial surface proteins (Pober, 2002; Pober and Sessa, 2007; Beguin et al., 2019). In this work we investigate whether *B. burgdorferi* leverages spirochete-induced activation of the vascular endothelium by establishing new interactions with the endothelium that were inaccessible prior to activation.

Bacterial surface proteins are the first point of contact between pathogens and the host milieu; those that demonstrate binding to host components are referred to as adhesins. *B. burgdorferi* contains approximately 85 surface-bound lipoproteins, primarily encoded by the large number of plasmids carried by *Borrelia* spirochetes (Dowdell et al., 2017). A number of characterized surface proteins display adhesin activity (see recent adhesin reviews (Caine and Coburn, 2015; Lin et al., 2017; Coburn et al., 2021; Stevenson and Brissette, 2023; Hejduk et al., 2025)). Binding occurs primarily to extracellular matrix components and proteins in the complement pathway (Barbosa and Isaac, 2018; Dulipati et al., 2020; Lin et al., 2020a; Skare and Garcia, 2020). However, it remains unknown whether any of these adhesins promote vascular interactions under shear stress. Addressing this question has been challenging because wild-type Lyme spirochetes encoding BBK32 and VlsE, display high levels of vascular adhesion that obscure binding contributions by other *B. burgdorferi* proteins. Recently, we identified VlsE as a potent vascular adhesin and generated a BBK32, VlsE doubly deficient strain that exhibits reduced vascular interactions (Tan et al., 2022), allowing its use as a starting point to investigate vascular interactions promoted by other adhesins.

In the work reported here, using intravital imaging, we show that *B. burgdorferi-*induced endothelial activation (Tan et al., 2021; Tan et al., 2023) generates new binding targets for *B. burgdorferi,* enabling previously unrecognized spirochete-vascular interactions under shear force. We identified five spirochete adhesins that specifically promote microvascular interactions only with activated endothelium, revealing a previously unknown pathogenic strategy for this important disease-causing organism. In addition, we report six new adhesin activities at play in early infection that do not require endothelial activation.

## Results

### Endothelial activation promotes new microvascular adhesion capabilities of *B. burgdorferi*

In our previous work, we demonstrated that *Cd1d^-/-^* mice exhibit endothelial activation in post-capillary venules following injection with *B.burgdorferi* and that this activation proceeded to potentiation, a term we use to describe endothelial permissiveness for spirochete extravasation at 24 h post-infection (Tan et al., 2021). Viewing transmigrated spirochetes at 24 h by intravital imaging (Kumar et al., 2015; Lin et al., 2020b; Tan et al., 2021; Tan et al., 2023) required the use of *Cd1d^-/-^* mice, which lack invariant natural killer T (*i*NKT) cells responsible for clearing *B. burgdorferi* in joint peripheral tissue (Lee and Hurwitz, 1992; Lee et al., 2010; Lee et al., 2014). However, the study of vascular adhesion directly after inoculation (Moriarty et al., 2008; Norman et al., 2008; Moriarty et al., 2012; Tan et al., 2022) does not require the use of Cd1d^-/-^ mice. Because wild-type (WT) mice are preferred to study host-pathogen interactions, including the effect of endothelial activation on spirochete adhesion, it was first necessary to establish whether activation of the endothelium occurred in WT mice following exposure to *B. burgdorferi.* To address this, we used BALB/c mice, which are the background of the *Cd1d^-/-^* mice used in our earlier studies on vascular transmigration.

**Fig. 1** shows the temporal dynamics of endothelial activation following injection of WT *B. burgdorferi* into BALB/c mice. Endothelial activation was assessed by quantifying adherent neutrophils in peripheral knee joint post-capillary venules using intravital microscopy (IVM). Representative images of anti-Ly6G–labeled (red) neutrophils in uninfected (t= 0) and infected mice (t = 3-20 h) are shown in **Fig. 1A** and **Video 1**. Quantitative analysis (**Fig. 1B**) revealed that in BALB/c mice endothelial activation was detected as early as 3 h post-infection and persisted through 20 h.

**Fig. 1.**
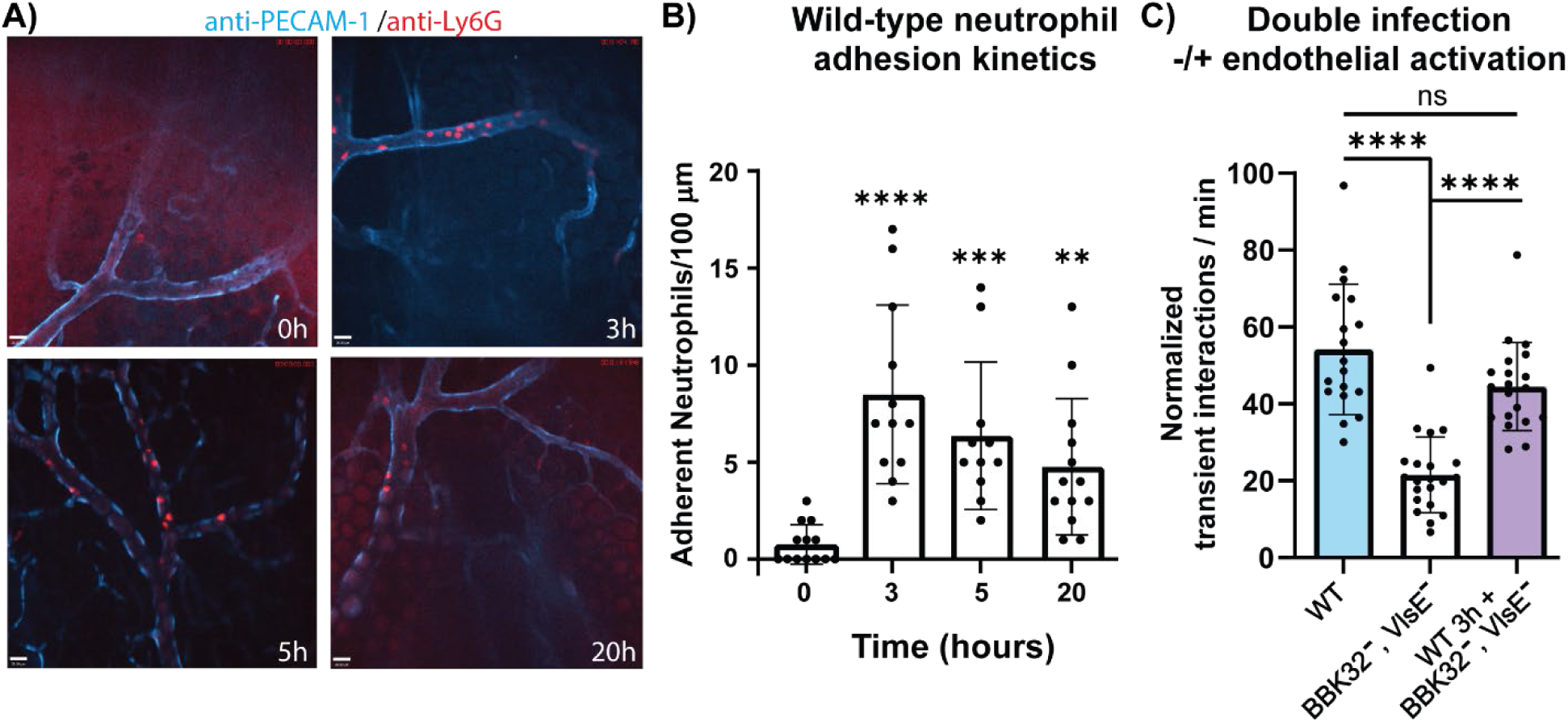
Endothelial activation in the knee joint-proximal tissue of WT *B. burgdorferi*-infected BALB/c mice. Neutrophil adhesion to the endothelium was assessed using intravital imaging. **A)** Representative micrographs show neutrophil recruitment at various times (3-20 hours) in BALB/c mice after infection with wild-type GFP-expressing *B. burgdorferi* (GCB726). Each group consisted of 3 mice, injected with 3 × 10^8^ spirochetes per mouse. Uninfected mice injected instead with phosphate-buffered saline (PBS) served as negative controls (T0). Neutrophils were stained with anti-Ly6G (red), and joint-proximal venules were stained with anti-PECAM-1 (blue), as described in the Materials and Methods section and previous studies. Scale bar = 20 µm. **B)** The graph represents total neutrophil adhesion after infection, defined as neutrophils that remained stationary for at least 30 seconds within a 100-μm length of unbranched venule (20 to 40 μm in diameter) over 3 minutes. Statistical significance was determined using the Kruskal-Wallis test with Dunn’s multiple comparison test. *p* < 0.05 was considered significant. Data represent the mean ± standard deviation (SD) from three independent experiments. **C)** The number of transient adhesion events per minute was quantified by intravital microscopy and normalized as described in Materials and Methods. Mice were injected with either a GFP-expressing wild-type strain (GCB726; n = 5) cultured with 1% blood (blue bar) and imaged 3 h post-injection; a GFP-expressing BBK32, VlsE doubly-deficient strain (GCB4038) cultured with 1% blood (white bar) and imaged 3 h post-injection; or subjected to a sequential double infection consisting of a non-fluorescent wild-type strain for 3 h followed by immediate imaging of the GFP-expressing BBK32, VlsE doubly-deficient strain (violet bar). The graph depicts the number of microvascular interactions observed in mouse venules. Statistical significance was assessed using the Kruskal–Wallis test with Dunn’s multiple-comparison post-hoc test. In all analyses, *p* < 0.05 was considered significant; \*\**p* < 0.01, \*\*\**p* < 0.001, \*\*\*\**p* < 0.0001, ns = not significant.

We then investigated whether endothelial activation resulted in any new shear force-resistant microvascular interactions with *B. burgdorferi.* This analysis was performed using i.v. inoculation with a doubly deficient GFP-expressing *B. burgdorferi* strain lacking both BBK32 and VlsE, which exhibits low levels of vascular interactions. Adhesion was assessed either immediately following infection (**Fig. 1 C,** white bar) or after endothelial activation induced by a prior injection of non-fluorescent WT *B. burgdorferi* administered 3 h before intravital imaging (**Fig. 1C**, violet bar). A GFP-expressing WT strain was included as a control for adhesion (**Fig. 1C**, light-blue bar). These data show that endothelial activation restores the adhesion capabilities of a doubly deficient *B. burgdorferi* strain to levels comparable to WT, suggesting the existence of previously undescribed adhesin(s) targeted to activated endothelium.

### Video S1

Intravital imaging of neutrophil adhesion to the endothelium in BALB/c mice following infection with wild-type GFP-expressing B. burgdorferi (GCB726). Neutrophils are labeled with anti-Ly6G (red) and venules with anti-PECAM-1 (blue). The left panel video shows the uninfected control (PBS-injected mice), the center panel is a representative video at 5 h, and the right panel is a representative video at 20 h post-infection. The videos display a 200-stack acquisition at 15 fps.

### Cytokine signaling promotes new microvascular adhesion capabilities of *B. burgdorferi*

Endothelial activation following exposure to *B. burgdorferi* results from cytokine signaling (Tan et al., 2021). Because the use of a non-fluorescent WT *B. burgdorferi* injection to activate the endothelium as performed in **Figure 1** was labor-intensive and time consuming for subsequent experiments, we sought to determine whether direct cytokine-induced endothelial activation in BALB/c mice could restore microvascular interactions of the doubly deficient BBK32^-^, VlsE^-^ strain (**Fig. 1**).

We selected a 3-hour exposure period for cytokine treatment and evaluated both known endothelial activators (GM-CSF and IL-1β) and potentiators (TNF-α, MCP-1 and IL-10) (Tan et al., 2021). All these cytokines restored the adhesion of the BBK32^-^, VlsE^-^ deficient strain (**Fig. 2**). A WT strain served as positive control (light blue bar) and the doubly deficient strain in the absence of cytokine treatment as a negative control (white bar). IL-17A, a cytokine with no known activator/potentiator activities, was also included as negative control (gray bar). Representative videos of WT spirochetes, the BBK32, VslE doubly deficient strain and the BBK32, VslE doubly deficient strain following MCP-1 treatment are shown in **Video S2**.

**Fig. 2.**
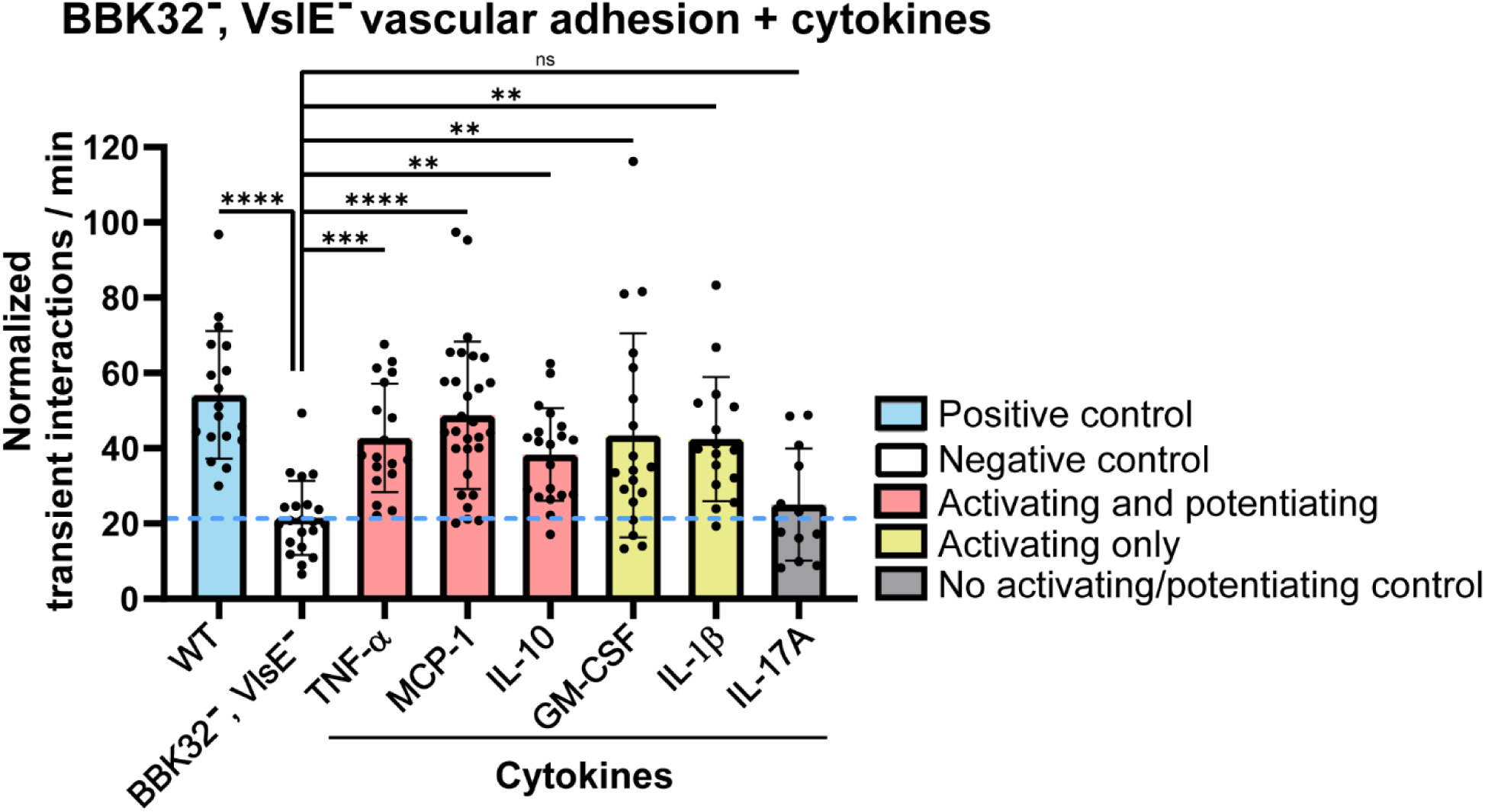
Effect of cytokines on microvascular interactions of the doubly deficient BBK32, VlsE strain of *B. burgdorferi* in BALB/c mice. Mice were injected with the wild-type strain (GCB726, n = 5) cultured with 1% blood (see Materials and Methods) for a positive control. For the analysis, they were injected with PBS (n = 5) or the specific cytokine 3 hours before infection with the GFP-expressing VlsE, BBK32 doubly deficient strain (GCB4036 or GCB4038) cultured with blood. Vascular interactions were detected using high acquisition rate spinning disk confocal intravital microscopy 5-50 minutes post-infection. The number of interactions per minute for transient adhesions was analyzed using intravital microscopy. The graph shows the number of microvascular interactions in mouse venules. Statistical significance was determined using the Kruskal-Wallis test with Dunn’s multiple comparison test. *p* < 0.05 was considered significant; \**p* < 0.05, \*\**p* < 0.01, \*\*\**p* < 0.001, \*\*\*\**p* < 0.0001, ns = not significant. The cytokines used were: TNF-α (45 ng/25 g mouse, n = 4), MCP-1 (5 μg/25 g mouse, n = 6), IL-10 (2.5 μg/25 g mouse, n = 5), IL-1β (250 ng/25 g mouse, n = 5), GM-CSF (15 ng/25 g mouse, n = 4) injected intravenously, and IL-17A (0.5 μg/25 g mouse, n = 3) injected intraperitoneally.

### Video S2

BALB/c mice were injected with the wild-type strain GCB726 (n = 5), cultured with 1% blood, as a positive control (left panel). For experimental analysis, mice were injected with phosphate-buffered saline (center panel) or MCP-1, 3 h prior to infection with the GFP-expressing VlsE/BBK32 doubly-deficient strain (GCB4036 or GCB4038) cultured with blood (right panel). Vascular interactions were visualized by high–acquisition rate spinning-disk confocal intravital microscopy 5–50 min post-infection. The videos correspond to a 200-stack acquisition at 15 frames per second (fps).

### Host adaptation is required to promote new microvascular adhesion capabilities of *B. burgdorferi* to activated endothelium

In the above experiments, the strain lacking the major adhesins BBK32 and VlsE showed restored binding to activated endothelium using our standard growth conditions for vascular adhesion experiments (Moriarty et al., 2008), which include growth with 1% blood to host-adapt the spirochetes. When the blood-exposure step was omitted, even in presence of MCP-1, the interaction of the spirochetes with the endothelium was not restored (**Fig. 3A**). This indicates that spirochete adaptation to the host environment is required to establish new interactions with activated endothelium and that it appears to be required to induce the synthesis of adhesin(s) involved in the interactions. Moreover, when we cultivated the strain in blood and subsequently treated the spirochetes with proteinase K, the adhesion was abolished (**Fig. 3B, Video S3**), indicating an involvement of a surface-exposed protein(s) in the interaction.

**Fig. 3.**
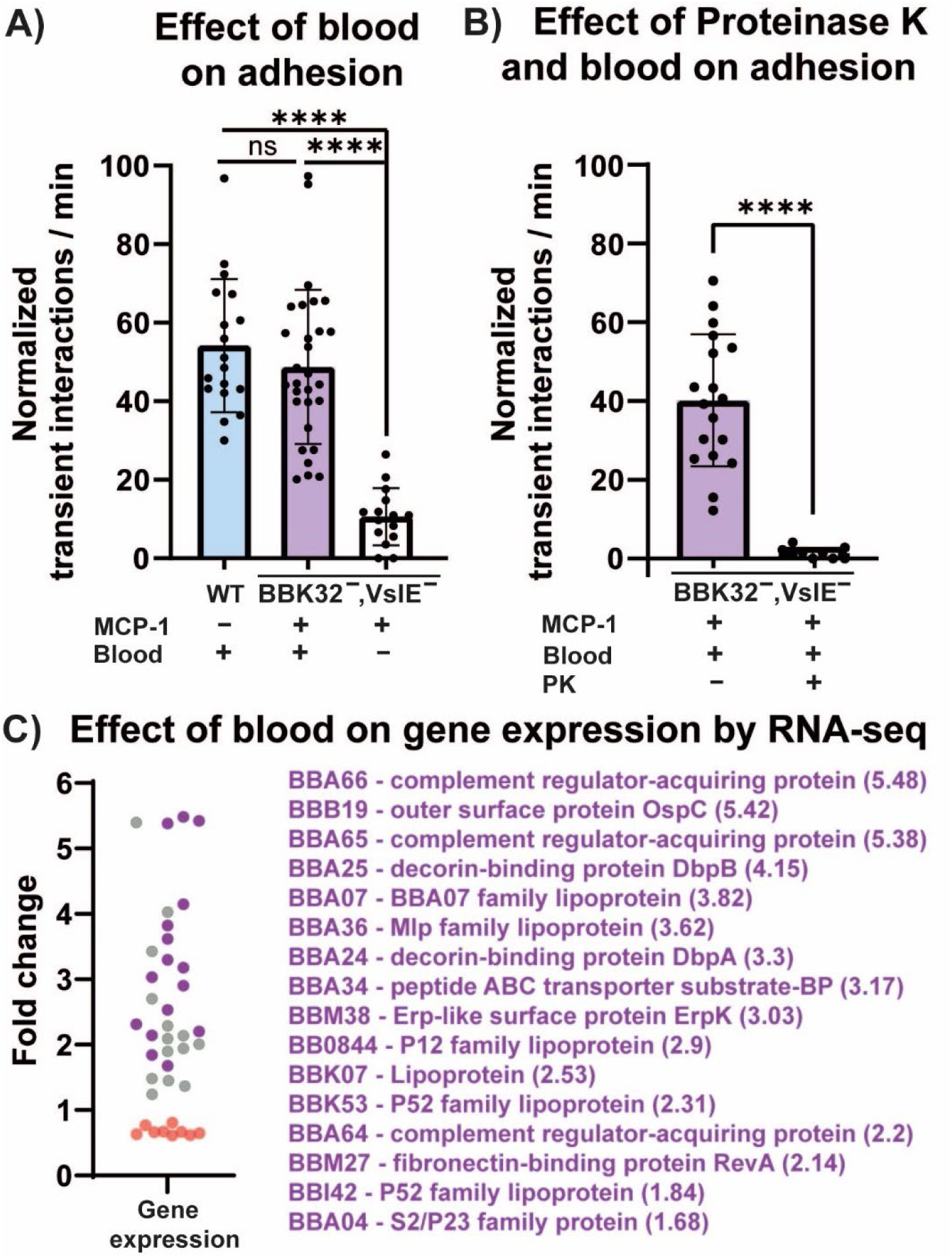
Effect of blood on the BBK32, VlsE minus strain of *B. burgdorferi*. **A)** Effect of blood exposure on microvascular interactions of the *Borrelia burgdorferi* doubly deficient strain in MCP-1–treated BALB/c mice. The GFP-expressing wild-type strain (GCB726), cultured with blood, was used as a positive control. GCB4038, which is deficient in both VlsE and BBK32, was grown with or without mouse blood for 48 hours prior to infection. To evaluate the impact of blood exposure on vascular adhesion, BALB/c mice (n = 3) were pretreated with 5 μg/25 g body weight of MCP-1 via intravenous injection 3 hours before infection. Vascular interactions were visualized by intravital microscopy as described in Fig. 1. Error bars represent standard deviation (SD). Statistical significance was assessed using the Kruskal–Wallis test followed by Dunn’s multiple comparisons; *p* < 0.05 was considered significant (\*\*\*\**p* < 0.0001, ns = not significant). **B)** Effect of proteinase K on microvascular transient interactions of the BBK32, VlsE minus strain of *B. burgdorferi* grown in blood in MCP-1 treated BALB/c mice. GFP-expressing doubly deficient strain GCB4036 was grown with mouse blood 48 hours before infection. Mice (n=3-4) received 5 μg/25 g mouse MCP-1 by i.v. injection 3 hr prior to infection with GCB4036. GCB 4036 was incubated with or without proteinase K 150 µg/ml one hour before infection. After infection, vascular transient adhesion interactions were detected by intravital microscopy as in Fig. 1. Error bars represent SD, and statistical significance was analyzed using the Mann-Whitney test; *p* < 0.05 was considered significant (\*\*\*\**p* < 0.0001). **C)** Differential gene expression in the doubly deficient *B. burgdorferi* GCB4036 following exposure to whole blood. Cultures were incubated with whole blood for 48 hours prior to RNA sequencing. A total of 39 genes were differentially expressed, including 30 upregulated genes (violet and grey dots) and 9 downregulated genes (red dots). Among the upregulated genes, 16 (shown in violet and listed on the right with the fold change values in parentheses) are predicted to encode surface-exposed proteins, based on the surface proteome (Dowdell et al., 2017).

To pursue identification of the blood-induced adhesin(s), we incubated the BBK32^-^, VlsE^-^ strain with 1% whole blood for 48 hours under the conditions used for the above intravital experiments and then performed RNA sequencing (**Fig. 3C**). A total of 39 genes were found to be differentially expressed, including 30 upregulated genes (violet and gray dots) and 9 downregulated genes (red dots). A previous study using 5% blood induction identified many more upregulated proteins (Tokarz et al., 2004), however, for this study it was necessary to use the same conditions as those used in our intravital studies and shown here in **Fig. 3A**. Among the upregulated genes we observed, 16 (shown in violet) and listed on the right with the fold change value in parentheses) are known or predicted to encode surface-exposed lipoproteins, based on multiple previous studies (see reviews by (Caine and Coburn, 2015; Lin et al., 2017; Coburn et al., 2021; Stevenson and Brissette, 2023; Hejduk et al., 2025)) and on the surface proteome defined by Dowdell and coworkers (Dowdell et al., 2017).

### Video S3

The GFP-expressing doubly-deficient strain GCB4036 was incubated in blood as described previously and either directly injected into MCP-1–treated mice (left panel) or preincubated with proteinase K (150 μg/ml) for 1 h prior to infection (right panel). Following infection, transient vascular adhesive interactions were visualized by intravital microscopy using the same acquisition and analysis conditions as in the preceding video. The videos correspond to a 200-stack acquisition rendered at 15 frames per second (fps).

### Identification of shear force-resistant adhesins

To identify the adhesins whose production was stimulated by growth in blood and that were capable of mediating shear force-resistant interactions, we cloned each candidate (**Fig 4A**) under the control of the medium-strength *resT* promoter (Bandy et al., 2014; Takacs et al., 2018) into an lp5-derived shuttle vector (Castellanos et al., 2018). This vector is maintained in linear form in *B. burgdorferi* and is compatible with pTM61, which carries the gene for GFP (**Table S1**). The resulting adhesin constructs were then introduced into GFP-expressing HB19 high passage (**Table S1**), a minimally adhesive strain that retains only four plasmids: cp26, lp17, lp54 and lp28-6 (Jacob Lemieux personal communication). Finally, mice were injected with each strain, and microvascular interactions in knee joint peripheral tissue were analyzed by intravital imaging (**Fig. 4A**). We analyzed the 16 upregulated predicted surface protein candidates, as well as BBK32 as a positive control (Norman et al., 2008; Moriarty et al., 2012) and P66 which we have previously studied but whose shear-resistant properties were unknown (Kumar et al., 2015; Tan et al., 2023). The data, shown in **Fig. 4B** revealed six new surface proteins (indicated in orange) with shear force-resistant adhesin activity: BB0844, BBA04, BBA65, BBK07, BBK53, ErpK (BBM38) and OspC, which displayed stable adhesion capabilities over the 50 min period analyzed (**Table S2**).

**Fig. 4.**
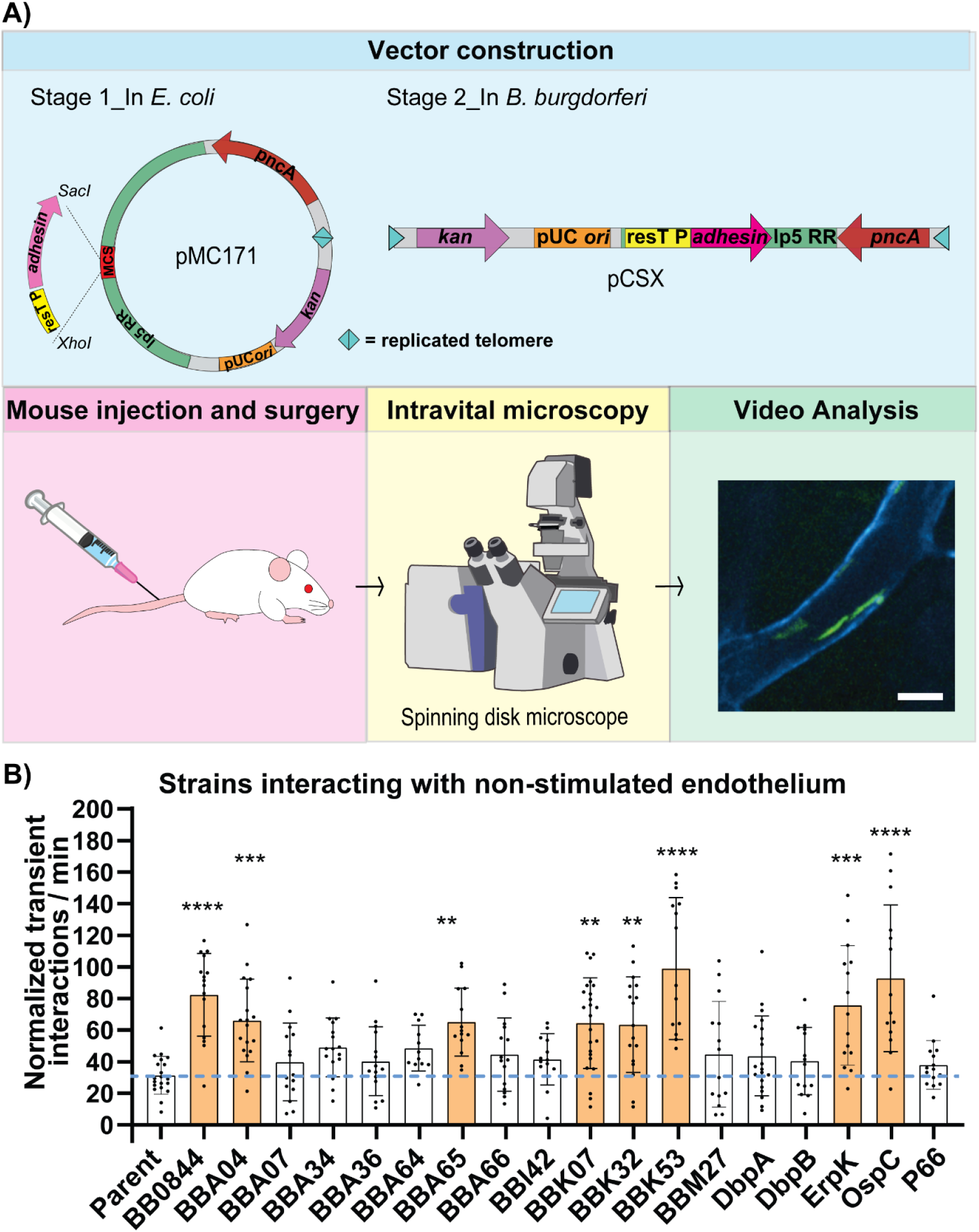
Identification of shear force-resistant adhesins binding to pre-activated endothelium. **A)** Schematic overview of the experimental workflow. Each selected surface-exposed protein was first cloned into a lp5-derived shuttle vector (stage 1, light blue box) and subsequently introduced into *B. burgdorferi* where the plasmid exists in linear form (stage 2, light blue box). The resulting strains were injected into the tail veins of BALB/c mice (pink box), after which surgical preparation and intravital imaging were performed (yellow box). Finally, the video analysis was performed as described in each case. Example of a micrograph, scale bar = 35 µm (green box). **B)** Quantification of transient interactions per minute. Spirochetes (3x10^8^) were introduced by tail vein injection into the parental strain (negative control, GCB989) or the indicated adhesin-producing strains (see **Table S1**). Transient vascular interactions within post-capillary venules were detected using high-acquisition-rate spinning disk confocal intravital microscopy in knee joint peripheral tissue 5-50 minutes post-infection. The number of normalized transient adhesion events per minute was plotted and analyzed. Error bars represent SD. Statistical significance was determined using the Kruskal-Wallis test followed by Dunn’s multiple comparisons; *P* < 0.05 was considered significant; \*\**p* < 0.01, \*\*\**p*< 0.001, \*\*\*\**p* < 0.0001, ns = not significant.

In our data analysis, we also noted that the size of the vessel where interactions were measured could have an effect for each adhesin; for some adhesins there was little effect, while for others the impact was greater. As the vessel size increases there are changes in the velocity of blood flow and shear stress but there can also be differences in protein content of the vascular bed as well as changes in the glycocalyx; therefore, a strict correlation between binding affinity in vessels of varying size is not possible. However, a simplistic assumption is that spirochete adhesion may be more difficult in larger vessels and that spirochetes carrying stronger adhesins would be less influenced by changes in vessel diameter, while those with weaker adhesins would display greater changes in binding. In an effort to predict a tentative relative binding strength of the various adhesins we calculated the Spearman correlation coefficient (r) for vessel size versus number of interactions per minute. The magnitude of r reflects the strength of the association: an r ≈ 0.1–0.3 indicates a weak relationship, r ≈ 0.3–0.6 indicates a moderate relationship, while r > 0.6 indicates a strong relationship. The predicted ranking of adhesion strength is presented in **Table 1**. A negative Spearman r indicates that adhesion decreases as vessel size increases.

**Table 1.**
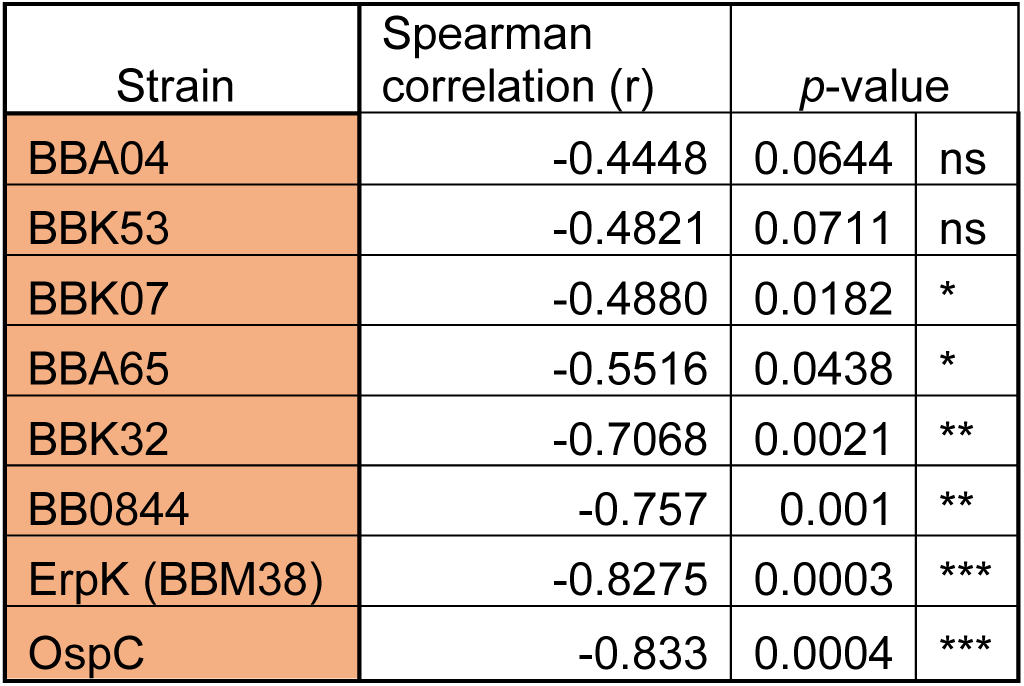
Predicted relative strength of the shear force-resistant adhesins.

Strains carrying BBA04 and BBK53 exhibited high levels of interactions per minute (**Fig. 4**) and consistently high adhesion across all vessel sizes, with no significant correlation between binding and vessel size, indicating that their attachment behaviour was independent of vessel geometry.

In contrast, strain BBK07 and BBA65 showed a moderate, but significant Spearman correlation, while BBK32, BB0844, ErpK (BBM38), and OspC, demonstrated progressively stronger size-dependent effects, as reflected by increasingly negative r and more statistically significant *p-*values. The ranking presented in this table should be considered theoretical at present and will require ex vivo studies for further investigation.

### Identification of shear force-resistant adhesins binding specifically to activated endothelium

We next compared vascular adhesion mediated by each of the above proteins in the presence versus the absence of MCP-1, shown in **Fig. 5**. Interestingly, the proteins fell into four distinct classes. The first class comprises five adhesins whose binding does not require endothelial activation by MCP-1 (**Fig. 5A**). The second and primary group of interest, includes proteins that did not display vascular interactions in the absence of MCP-1 but did display interactions with activated endothelium (**Fig. 5B**). This novel class of adhesins was composed of BBA07, BBA36, BBA66, DbpA, and P66. Surprisingly, we also identified a third class of adhesins which demonstrate reduced vascular interactions (**Fig. 5C**) following MCP-1 promoted endothelial activation (BBK53, OspC and ErpK (BBM38)). We propose that this reduction may represent a coordinated **’**attachment-detachment’ switch necessary for dissemination which we elaborate on in the Discussion section. Finally, six of the proteins assayed did not display adhesin activity with either activated or pre-activated endothelium (**Fig. 5D**).

**Fig. 5.**
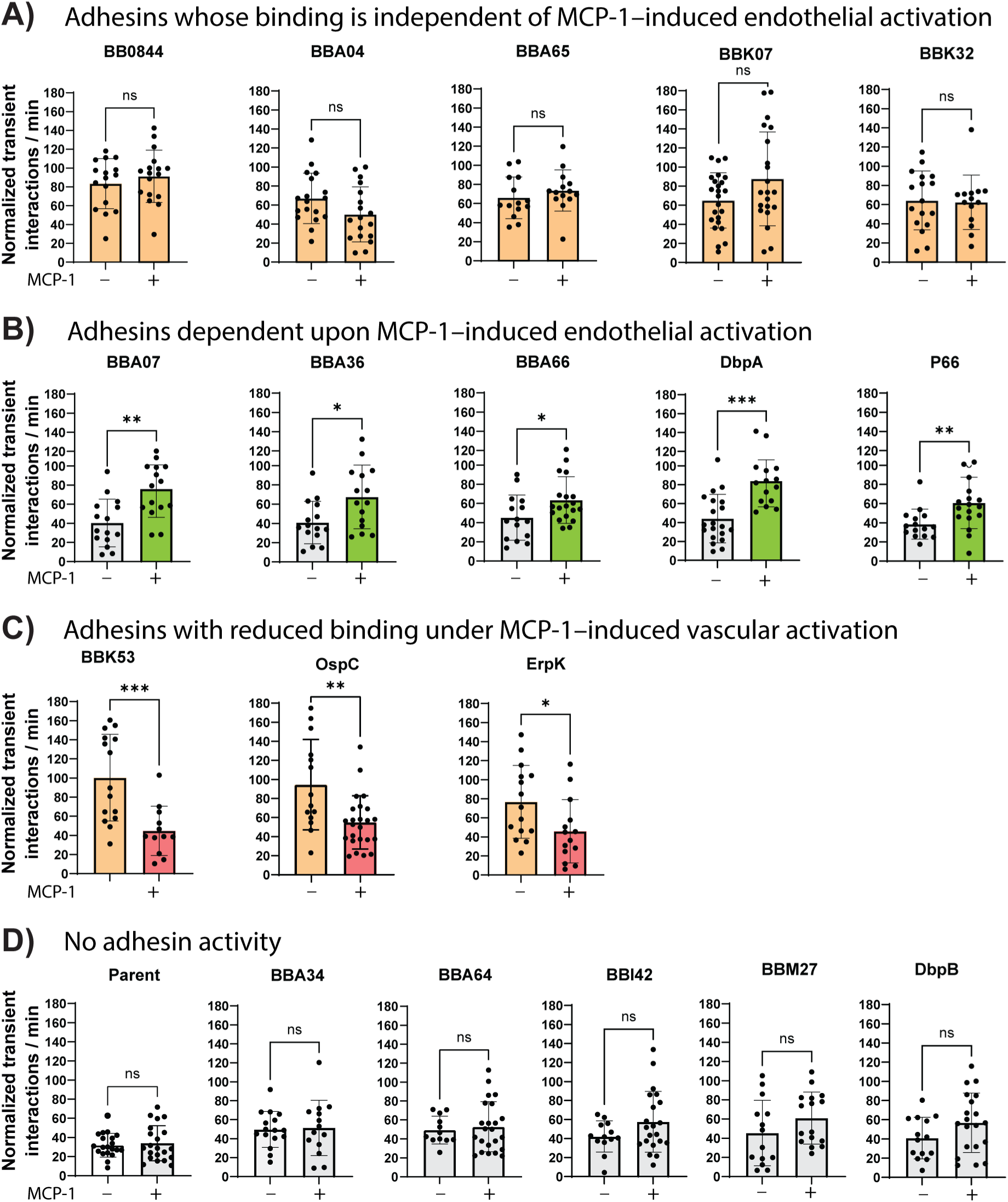
Effect of MCP-1 on microvascular interactions in BALB/c mice. Mice were injected with PBS or MCP-1 (5 µg/25g mouse) 3 hours before infection with strains carrying the indicated surface-exposed proteins. Vascular adhesion interactions were detected using high acquisition rate spinning disk confocal intravital microscopy 5-50 minutes post-infection. The number of interactions per minute for transient adhesions was analyzed. The graph shows the comparison of the number of microvascular interactions with or without MCP-1 in mouse venules for each protein. Statistical significance was determined using the Mann-Whitney test with for each strain comparison (with/without MCP-1). In all cases P < 0.05 was considered significant, \**p* < 0.05, \*\**p* < 0.01, *** *p* < 0.001, **** *p* < 0.0001, ns = not significant.

Next, we also assessed the relative strength of MCP-1 dependent adhesins by calculating the Spearman correlation coefficient (r) between number of adhesions and vessel diameter. Strains BBA66 and strain P66 exhibited consistently high adhesion across all vessel sizes, and showed weak, non-significant Spearman correlation, indicating that their attachment was independent of vessel size. In contrast, strains BBA36 and BBA07 showed a moderate, but statistically significant Spearman correlation, whereas DbpA showed a stronger, size-dependent statistically significant Spearman correlation. Based on these analyses, the adhesins were tentatively ranked by relative adhesion strength, summarized in **Table 2**. The relative strength is in descending order: BBA66>P66>BBA36>BBA07>DbpA.

**Table 2.** Relative strength of MCP-1 dependent adhesins.

| Strain | Spearman correlation ($r$) | $p$-value | |
| --- | --- | --- | --- |
| BBA66 | -0.2334 | 0.3361 | ns |
| P66 | -0.3168 | 0.2002 | ns |
| BBA36 | -0.5286 | 0.0454 | * |
| BBA07 | -0.5821 | 0.0252 | * |
| DbpA | -0.6393 | 0.0122 | * |

## Discussion

In the ever-escalating battle between pathogens and hosts, intricate strategies and counter-strategies are the evolutionary norm. This work reveals a previously unknown tactic in the arsenal of the Lyme disease bacterium. Shortly after infection, signals from *B. burgdorferi* promote endothelial activation through the action of P66, a surface protein that induces cytokine release primarily from host neutrophils (Tan et al., 2021). It has been long-established that during transmission/early infection, signals from the host result in changes in gene expression and adapt the spirochete for the infection process (Iyer and Schwartz, 2016; Samuels et al., 2021). In the current work, we show that host adaptation by the spirochetes results in a series of new shear force-resistant *B. burgdorferi*-endothelial interactions that are targeted only to activated endothelium. We have identified five spirochete-encoded adhesins responsible for this previously undescribed activity (blue box, center **Fig. 6**). Interestingly, two of these adhesins (P66 and DbpA) have been previously shown (Tan et al., 2023) to serve as endothelial potentiators (see **Fig. 6**), a function which occurs after endothelial activation and which might be expected to be facilitated by enhanced endothelial binding. P66 also plays a role in the subsequent extravasation event (Tan et al., 2023). The specific binding to activated endothelium of these two adhesins is therefore correlated with their choreographed function in vascular transmigration.

**Figure 6.**
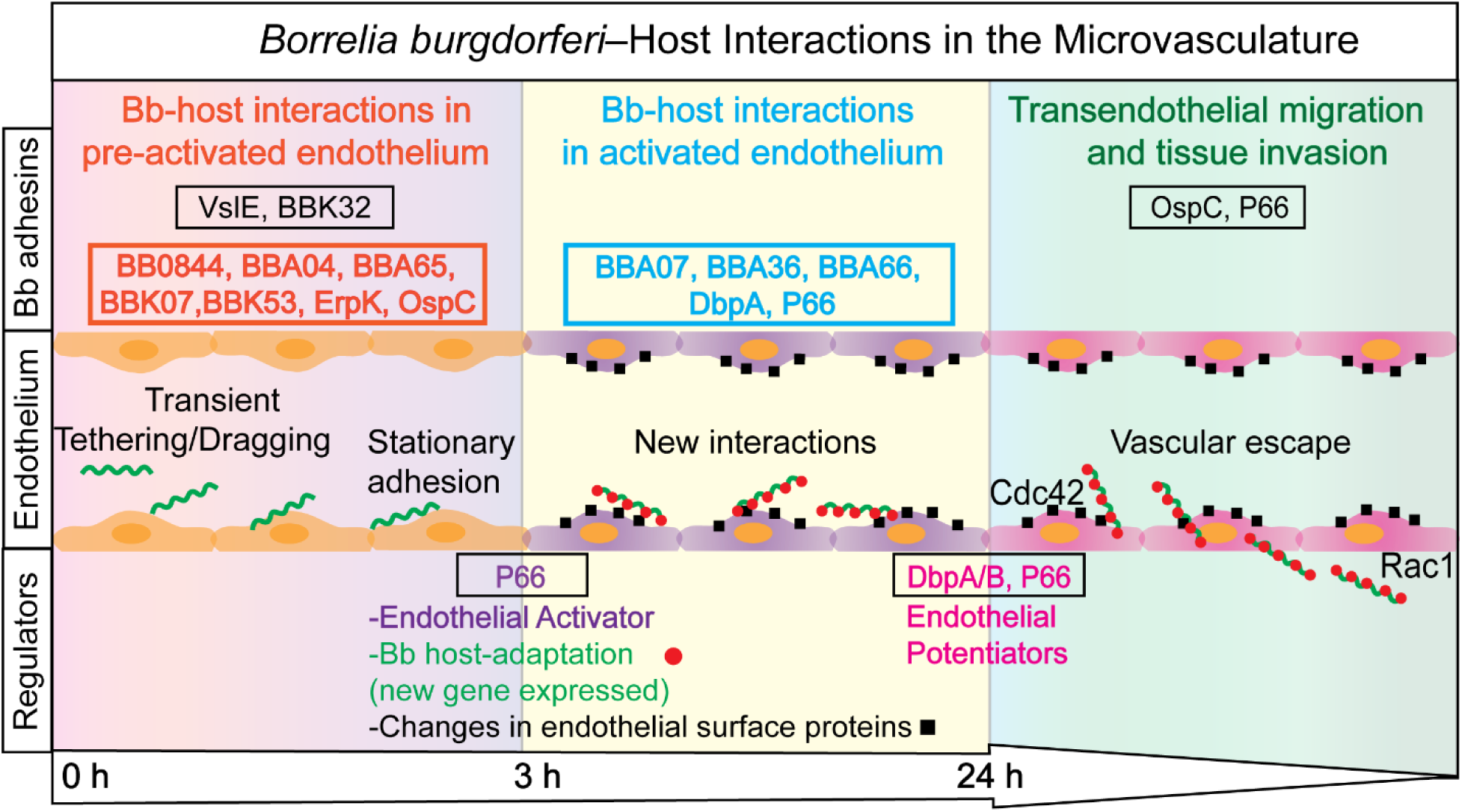
Schematic of the role of *B. burgdorferi* adhesins in hematogenous dissemination. The schematic shows a sequential adhesion cascade: initial tethering (red box) is followed by inflammation and engagement of activation-dependent adhesins (blue box) to facilitate vascular escape. In pre-activated endothelium (left panel) there are two known shear force-resistant adhesins: VlsE, and BBK32 and BB0844, BBA04, BBA65, BBK07, BBK53, ErpK (BBM38) and OspC described here (red box). As part of host-adaptation (middle panel), spirochetes in contact with blood express other adhesins, that show adhesion only when endothelium is activated: BBA017, BBA36, BBA66, DbpA and P66 (blue box). Vascular escape (right panel) requires the activity of OspC and P66 as previously described. For example, P66 is involved in activating and potentiating the endothelium and in vascular escape; these activities require its integrin binding residues (Tan et al., 2021; Tan et al., 2023) (Kumar et al., 2015). DbpA/B serve to potentiate the endothelium (Tan et al., 2023) and OspC functions in vascular escape (Lin et al., 2020b; Tan et al., 2023). Host proteins Cdc42 and Rac1 also appear to play a role in transendothelial migration (Alvarez-Olmedo et al., 2025).

Interestingly, in this study we also observed that endothelial activation (via MCP-1) significantly reduces vascular interactions mediated by BBK53, OspC, and ErpK (**Fig. 5C**). This observation may have different explanations. Reduced endothelial interactions following activation could result from a loss of the ligands for each adhesin upon endothelial activation. Two of the three adhesins (OspC and ErpK) are known glycosaminoglycan (GAG) binders (Brissette et al., 2008; Lin et al., 2020b). Endothelial activation induces profound remodeling of GAGs in both the glycocalyx and the extracellular matrix (Witjas et al., 2019; Dogne and Flamion, 2020; Moore et al., 2021; Li et al., 2024), as well as changes in endothelial surface protein composition (Beguin et al., 2019). Alternatively, new endothelial surface proteins may obscure access to the ligands for these adhesins.

Regardless of the mechanism, this reduction in binding might represent a coordinated ‘attachment-detachment’ switch necessary for dissemination whereby these adhesins might function as ‘tethering anchors’ on pre-activated endothelium. Upon the successful induction of local inflammation, signaled by cytokine release, the spirochetes would disengage from luminal receptors so they can transmigrate the vessel. Thus, the downregulation of these specific interactions may not represent a simple loss of function, but may serve as a prerequisite for the subsequent engagement of activation-dependent adhesins (e.g., P66, DbpA) and final transendothelial migration. Such a biphasic adhesion strategy would be similar to the dynamic binding behavior of leukocytes during recruitment and transmigration (McEver and Zhu, 2010; Sundd et al., 2013).

During the course of these studies, we also uncovered seven new spirochete adhesins that are involved in previously undescribed shear force-resistant interactions with the endothelium before it becomes activated (red box, left **Fig. 6**). This dramatically increases the number of known adhesins to preactivated endothelium to eight with more no doubt remaining to be discovered. A possible role for these adhesins may be to aid in dissemination by directing spirochetes to a variety of different tissue types. One of these adhesins, OspC has been previously implicated in joint invasion and colonization (Lin et al., 2020b) as well as in spirochete extravasation into the knee joint (Tan et al., 2023). This role may require a reduced binding affinity to that observed early in infection (**Fig. 6**).

### The methodology

In this work we used intravital microscopy to visualize *B. burgdorferi-*endothelial interactions in living BALB/c mice in post-capillary venules in real time (Chaconas et al., 2020). While this approach uses i.v. injection of high numbers of spirochetes and deviates from the normal tick-mediated infection process, it is nonetheless a powerful methodology to study mechanistic issues related to hematogenous dissemination (Moriarty et al., 2008; Norman et al., 2008; Lee et al., 2010; Moriarty et al., 2012; Lee et al., 2014; Kumar et al., 2015; Kao et al., 2017; Lin et al., 2020b; Tan et al., 2021; Tan et al., 2022; Tan et al., 2023). We identified potential adhesins that were upregulated by growth in blood. These were cloned under the control of the medium-strength *resT* promoter and the resulting plasmids were used to transform *B. burgdorferi* HB19 high passage, which has low inherent adhesin activity. The resulting strains were injected into mice and microvascular interactions were scored in knee joint-proximal tissue in the presence and absence of MCP-1 to activate the endothelium. We previously used the general approach of introducing an exogenous gene to identify and verify BBK32 (Norman et al., 2008) and VlsE (Tan et al., 2022) as shear force-resistant adhesins. Here, this approach was successful in identifying five endothelial activation-dependent shear force-resistant adhesins as well as six previously uncharacterized activation-independent shear force-resistant adhesins. It should be noted, however, that many more adhesins of both types may exist but may have been missed by the narrow criteria used to focus the current study towards a successful outcome. Another limitation in our approach is that our intravital imaging was performed in the mouse knee joint peripheral tissue and based upon molecular variation in different vascular beds (Aird, 2007a; b), the results found in joint tissue may not be extrapolatable to the vasculature in other tissues.

Nonetheless, the approach reported here has identified endothelial activation-dependent adhesins, opening a new conceptual stage in the dissemination pathway and revealing a need for further studies using alternative approaches to further investigate this new area and identify additional such adhesins if they exist.

### The adhesins

Dissemination of the Lyme disease spirochete is important for disease pathogenesis. During hematogenous dissemination, *B. burgdorferi* utilizes an array of adhesins to facilitate its binding to and migration across endothelial cells to escape the vasculature and invade surrounding tissue. Before this study only two shear force-resistant adhesins of *B. burgdorferi* had been reported: BBK32 (Norman et al., 2008; Moriarty et al., 2012; Ebady et al., 2016; Niddam et al., 2017) and VlsE (Tan et al., 2022). Other adhesins were reported to play a role in dissemination (see **Fig. 6)**, but endothelial interactions withstanding shear force were not possible to test for in the presence of the strong adhesins BBK32 and VlsE. The use of a doubly deficient BBK32^-^ ,VlsE^-^ strain reduced endothelial interactions (Tan et al., 2022) and allowed the first observation of endothelial activation-requiring adhesin activity (**Fig. 1**), which was followed by functional gain experiments in high passage HB19, a strain that produces neither BBK32 nor VlsE. These experiments identified the endothelial activation-dependent and independent adhesins characterized here (**Fig. 6**, red and blue boxes). Although HB19 high passage encodes many adhesins, particularly those on lp54, the low intrinsic binding activity of this spirochete to post-capillary venules indicates that expression levels are low and any expression did not interfere with the identification of a number of adhesins present on exogenously added plasmids.

### Adhesins targeting activated endothelium

The five adhesins identified in order of proposed binding strength (**Table 2**) are BBA66>P66>BBA36>BBA07>DbpA. A summary of the properties of each of these adhesins is provided in the **Supplementary** section. It is noteworthy that except for P66, the other activation-dependent adhesins are carried on lp54 (Plasmid A), a highly conserved linear plasmid which is found in all *B. burgdorferi* isolates and is part of the core genome (Laing et al., 2025). The reader is referred to the **Supplementary** section and recent reviews for more information on the known *B. burgdorferi* adhesins of various types (Coburn et al., 2013; Brissette and Gaultney, 2014; Petzke and Schwartz, 2015; Hyde, 2017; Lin et al., 2017; Coburn et al., 2021; Stevenson and Brissette, 2023; Hejduk et al., 2025).

### Adhesins targeting endothelium before activation

The six new shear force-resistant adhesins that bind to pre-activated endothelium are BBA04>BBK53>BBK07>BBA65>BBK32>BB0844>ErpK>OspC, in order of proposed binding strength (**Table 2**). A summary of their properties is provided in the **Supplementary** section.

### Adhesin targets on activated endothelium

An interesting question raised by the results presented here is, what are the targets for *B. burgdorferi* binding that are specific for activated endothelium? Endothelial cells undergo profound phenotypic and molecular changes in response to inflammatory stimuli through the NF-κB signalling pathway, generating a surface landscape that differs markedly from that of resting endothelium and provides surface adhesion molecules including E-selectin, P-selectin, ICAM-1, VCAM-1 for leukocyte recruitment (Pober, 2002; Pober and Sessa, 2007). More recent studies have revealed the breadth and magnitude of endothelial activation induced by cytokines. Quantitative proteomic analyses show that approximately 136 endothelial proteins are differentially expressed including >30 cell surface proteins (Beguin et al., 2019). Adhesion receptors are among the most strongly induced, showing 20- to 32-fold increases in abundance. In addition to adhesion molecules, other induced proteins indicate that endothelial activation involves extensive remodeling of metabolic, redox, and extracellular matrix-associated pathways (Beguin et al., 2019), as well as post-translational modifications including phosphorylation and dephosphorylation of signaling and structural proteins. Such post-translational modifications may influence ligand availability, receptor clustering, or binding affinity, thereby further shaping the endothelial surface presented to circulating pathogens. The wealth of endothelial changes upon activation provides many new possible ligands for *B. burgdorferi* to target. Further studies into this new area will no doubt reveal fascinating interactions in the host-pathogen battle for supremacy and mechanistic roles for *B. burgdorferi* adhesins in endothelial transmigration, which allows subsequent tissue colonization and a wide variety of clinical manifestations.

## MATERIALS AND METHODS

### Ethical Approval and Use of Experimental Mice

All studies involving animals were conducted in accordance with the most recent guidelines and policies set forth by the Canadian Council on Animal Care, outlined in the Guide to the Care and Use of Experimental Animals. The protocol utilized in this research was approved by the Animal Care Committee at the University of Calgary (approval number AC24-0136-1). Wild-type BALB/c mice were acquired from Charles River Laboratories in Wilmington, MA. Both male and female mice aged 6 to 9 weeks were included in the experiments.

### Antibodies and reagents

Alexa Fluor 647–conjugated anti-mouse CD31 (PECAM-1; clone 390, cat. 102416) and phycoerythrin (PE)–conjugated anti-mouse Ly6G (clone 1A8, Rat IgG2 a, κ isotype, cat. 127608) were obtained from BioLegend Inc. (San Diego, CA). The recombinant mouse cytokines used in this study were all carrier-free and sourced from BioLegend. TNF-α (cat. 575202) was administered intravenously at a dose of 45 ng per 25 g mouse, and CCL2/MCP-1 (cat. 578404) was administered intravenously at 5 μg per 25 g mouse. IL-10 (cat. 575802) was delivered at 2.5 μg per 25 g mouse, whereas IL-1β (cat. 575102) was administered intravenously at 250 ng per 25 g mouse. GM-CSF was administered intravenously at 15 ng per 25 g mouse, and IL-17A (cat. 576002) was injected intraperitoneally at 0.5 μg per 25 g mouse.

### Plasmid construction: Cloning of Adhesin Genes Under the *resT* Promoter

To generate expression plasmids encoding the adhesins of interest, the mid-strength promoter *resT* and each adhesin gene were amplified from *B. Burgdorferi* strain 5A4 by PCR using the Phusion High-Fidelity DNA Polymerase kit (New England Biolabs). PCR amplifications were performed for 30 cycles of 95 °C for 30 s, 50 °C for 30 s, and 68 °C for 1 min, followed by a final extension at 68 °C for 7 min. Primers were designed to include 20 bp overlaps at the 5′ and 3′ ends of the genes to facilitate cloning (primer sequences in **Table S3**). Amplicons were purified using the QIAquick PCR Purification Kit (Qiagen), and DNA concentrations were measured with the Qubit dsDNA Kit (Thermo Fisher Scientific). The *resT* and adhesin fragments were assembled into the XhoI/SacI-digested pMC171 vector using the NEBuilder HiFi DNA Assembly Master Mix (New England Biolabs), following the manufacturer’s instructions. Ligation reactions were transformed into chemically competent *E. coli* DH5α, and kanamycin-resistant colonies were screened by using the PCR condition described using the same primer sets to verify the presence of both *resT* and the adhesin insert. Positive clones were submitted to Plasmidsaurus for nanopore plasmid sequencing (https://plasmidsaurus.com/). A list of the *E. coli* constructed strains can be found in **Table S4**.

### *B. burgdorferi* strain construction

Electrocompetent cell of the high passage HB19 strain (GCB 989) were prepared and transformed with 0.5-1 µg of the adhesin-encoding plasmid DNA as previously described (Samuels DS., 1995). Immediately following transformation, the cell-DNA mixture was transferred to 10 ml of pre-warmed BSK II supplemented 6% rabbit serum (PelFreez, cat 31125 or 31123). The transformation cultures were allowed to recover for 24 hrs. at standard culture conditions: 35 °C with 1.5 % CO_2_.

Following recovery, transformed cultures were expanded in additional BSK II media to a final volume of 100 ml and subsequently maintained under selective pressure with 200 µg/ml gentamycin and 100 µg/ml kanamycin. Afterwards, 250 µl aliquots of the culture were plated into 96-well plates for clonal isolation at 35 °C with 1.5 % CO_2_ until a characteristic medium color change (from red to yellow) indicated growth, typically within 7-10 days.

Wells that exhibited a characteristic medium color change (yellow), were screened by PCR to identify the presence of the kanamycin resistance gene using primers B70 and B71. PCR-positive transformants were sub-cultured further for gDNA extraction, which was done using the Puregene Cell Core Kit (Qiagen), according to the manufacturer’s instructions. Purified genomic-DNA preparations were subjected to further PCR analyses with the previously described conditions to verify the insertion of the adhesin construct and the products were evaluated by 0.5% agarose gel electrophoresis to confirm the expected plasmid sizes. A list of the *B.burgdorferi* constructed strains can be found in **Table S1**.

### Bacterial Strains and Culture Conditions

The B. burgdorferi strains employed in this study are listed in **Table S1** of the Supplemental Material. Spirochetes were cultured in Barbour-Stoenner-Kelly II (BSK-II) medium after inoculation from frozen glycerol stocks. Cultures were incubated at 35°C until reaching approximately 1-5 × 10⁷ cells/ml in 6 ml of BSK-II medium for regular maintenance. For intravital experiments, the spirochetes were cultured in 50 ml BSK-II medium for intravital experiments when required, 24-48 h prior the experiment 1% of total blood and triple antibiotic cocktail to a concentration of 0.02 mg/ml phosphomycin disodium salt cat. P5396, Sigma/ 0.0025 Amphotericin B from Streptomyces sp. cat. A4888, Sigma/ 0.05 rifampicin cat. R3501-5G, Sigma was added. Spirochetes were then collected by centrifugation at 6,000 × g for 15 min at 4°C, washed twice with 100 mL of cold PBS as described previously (30), and resuspended in cold PBS at a final density of 2 × 10⁹ cells/mL for mouse infections. E. coli strains were grown at 37°C in Luria-Bertani broth or on LB agar (BD Bioscience, Franklin Lakes, NJ), with or without ampicillin (100 μg/mL) as required.

### Mouse Infection and Intravital Imaging Procedures

For intravital microscopy, mice were anesthetized via intraperitoneal injection of ketamine (200 mg/kg; Bimeda-MTC Animal Health Inc., Cambridge, ON) and xylazine (10 mg/kg; Bayer Inc., Toronto, ON). Surgical preparation for imaging was performed as previously described (16, 30, 31). The imaging area was stabilized under a glass coverslip (VWR Scientific; cat. 48393-026; 22 × 30 mm) and visualized using a spinning-disk confocal microscope (Quorum Spinning Disk coupled with sCMOS camera: Photometrics Prime BSI (1966x1966) [95% QE, back illuminated, 80fps]). Blue (Ex 360/20nm, FT 400nm, Em 425nm) and Green lasers (Ex 470/40nm, FT 510nm, Em 515nm) were used.

### Neutrophil recruitment assay

Intravital microscopy was performed at the time points indicated in the figure legends, and neutrophil adhesion was quantified following established protocols (16). Blood vessels were labeled with anti-PECAM-1, and neutrophils were stained with PE-conjugated anti-mouse Ly6G. Adherent neutrophils were manually quantified offline using Volocity software (version 6.0.1). A cell was considered adherent if stationary for ≥30 s. Neutrophil adhesion was expressed as the number of adherent cells within a 100-µm segment of an unbranched venule (25–40 µm diameter) over a 5-min recording period.

### Assessment of *B. burgdorferi* adhesion

To evaluate whether cytokine treatment could rescue the vascular adhesion defect of the *vlsE^−^; bbk32*^−^ adhesin mutants, BALB/c mice were anesthetized with isoflurane and administered the appropriate cytokine 3 h before intravenous (tail vein) injection of 3 × 10⁸ GFP-expressing B. burgdorferi in 150 µL of PBS.

For quantification of spirochete interactions, 5–7 videos of vessels ranging from 20–40 µm in diameter were recorded for 3 min each. A 100-µm region of each vessel was analyzed, and all visible spirochetes passing through the area were counted. To account for differences in vessel size, the cross-sectional area of each vessel was calculated using π x r² and normalized to that of a 20-µm-diameter vessel to obtain a normalization factor (NF). Normalized transient interactions per minute were calculated as total adhesion events per minute ÷ NF.

### RNAseq

For RNAseq GCB4036 was grown in 50 ml BSK-II + 6% rabbit serum as the strain was grown for intravital microscopy in the early stages of this research. The culture was grown to a density of ∼5 x10^5^ and split in two. Fresh mouse blood was added to one of the two cultures to a final concentration of 1%. Each of the two cultures was then split into three 7 ml replicates and incubated at 35 degrees C for 48 hours at which point the cultures had reached a density of 3.3-4.3 x 10^7^ spirochetes/ml. A total of 1.32 x 10^8^ spirochetes for each replicate was harvested by centrifugation at 6,000xg at 4 degrees C for 15 minutes. To each pellet 0.5 ml TRIzol reagent (ThermoFisher, 15596026) was added and total RNA was purified using the PureLink RNA Mini Kit (Ambion, 12183020) including the On-column PureLink DNAse Treatment step. RNA integrity and concentration was checked for each replicate using TapeStation analysis (Agilent Technologies). Samples were submitted to the Centre for Health Genomics and Informatics, Cumming School of Medicine, University of Calgary for rRNA depletion, library preparation and sequencing using MiSeq V3 with 150 cycles.

## Acknowledgments

We would like to thank Genevieve Chaconas for technical support and John Leong and Jenifer Coburn for helpful discussions and comments on the manuscript. Intravital imaging was performed in the Live Cell Imaging Laboratory in the Snyder Institute for Chronic Diseases at the University of Calgary and DNA sequencing by the Centre for Health Genomics and Informatics, Cumming School of Medicine, University of Calgary. Research reported in this publication was supported by grants PJT-153336 from the Canadian Institutes of Health Research to George Chaconas and by a CIHR Postdoctoral Fellowship to Daiana Alvarez Olmedo and by the National Institute of Allergy and Infectious Diseases of the National Institutes of Health under award number R01AI169724.

## Notes

### Competing Interest Statement

The authors have declared no competing interest.

