## Supplementary material for "Making the best of a sticky situation: Infection-mediated endothelial activation promotes new interactions with adhesins of the host-adapted Lyme disease spirochete"

**Supplementary Material for Olmedo et al.**

### Description of adhesins binding specifically to activated endothelium

#### BBA66

BBA66 is an RpoS-regulated lipoprotein that is preferentially expressed under mammalian host–like conditions, particularly during the nymphal tick blood meal (Caimano et al., 2007; Patton et al., 2013). Genetic inactivation of *bba66* results in significant attenuation of infectivity during tick-mediated transmission, but not following needle inoculation (Maruskova and Seshu, 2008; Patton et al., 2013). Specific ligands for BBA66 remain to be defined, however it may be involved in close host–pathogen interactions, including colocalization with the host cytoskeletal adaptor Daam1 and promoting coiling phagocytosis in neuroglial cells (Antonara et al., 2007; Williams et al., 2018). Binding to activated endothelium adds a new function to those described for this protein.

### P66

Although P66 was not among the blood-induced proteins that we identified by RNA seq (**Fig. 3**) we nonetheless added it to our list of proteins to analyze due to its importance in the hematogenous dissemination pathway (**Fig. 6**). P66 is a conserved outer membrane protein that functions as both a porin (Skare et al., 1997; Pinne et al., 2007; Barcena-Uribarri et al., 2010; Barcena-Uribarri et al., 2013) and a β3-integrin–binding adhesin, positioning it as a central mediator of vascular interactions during hematogenous dissemination (Coburn et al., 1999; Coburn and Cugini, 2003; Antonara et al., 2007). It is the only integral membrane protein identified in our screen and the protein displaying the greatest number of functions in the hematogenous dissemination pathway (**Fig. 6**). P66 directly engages αvβ3 integrin on endothelial cells and its integrin binding residues promote endothelial activation, endothelial potentiation and transendothelial migration (Kumar et al., 2015; Tan et al., 2021; Tan et al., 2023). These intravital findings correlate with reduced dissemination of a P66 integrin binding mutant to secondary tissues, including the heart and joints, and highlight how P66-mediated integrin engagement coordinates adhesion, endothelial signaling, and vascular escape during early infection (LaFrance et al., 2011; Caine and Coburn, 2015; Ristow et al., 2015; Tal et al., 2024). Binding to activated endothelium adds a new function to the list for this important and versatile protein.

**BBA36**

BBA36 is a surface-exposed lipoprotein (Brooks et al., 2006; Dowdell et al., 2017). Transcriptional profiling shows *bba36* expression is regulated as part of the broader RpoS-dependent adaptive response (Caimano et al., 2007; Iyer et al., 2015). Although the specific host ligand(s) mediating BBA36-dependent interactions remain undefined, our data uncover a previously unrecognized function for BBA36 as a contributor to spirochete adhesion under conditions of endothelial activation, highlighting a role in host–pathogen interactions relevant to dissemination.

**BBA07**

BBA07 is a surface-exposed lipoprotein and is preferentially expressed during the nymphal tick blood meal as part of the RpoS-regulated gene program associated with mammalian infection (Caimano et al., 2007; Xu et al., 2010). Functional studies demonstrate that loss of *bba07* results in a pronounced defect in tick-to-mammal transmission but no defect in infection by needle inoculation (Xu et al., 2010). BBA07 has not previously been assigned a defined host ligand and has not been implicated in hematogenous dissemination. However, its transmission-associated expression profile places it within the cohort of surface proteins that facilitate host–pathogen interactions during early infection. Here, we identify a previously unrecognized contribution of BBA07 to spirochete adhesion in activated endothelium.

**DbpA**

The decorin-binding protein DbpA, whose expression is regulated by RpoS (Caimano et al., 2007) is a well-established surface adhesin that mediates high-affinity binding to decorin and decorin-associated collagen fibrils within the extracellular matrix, thereby promoting bacterial retention and persistence in the skin at the site of tick inoculation (Brown et al., 2001; Weening et al., 2008). Loss of DbpA and DbpB results in impaired dermal colonization and reduced infectivity (Blevins et al., 2008; Hyde et al., 2011) and plays a role in tissue tropism (Lin et al., 2014). Combined deletion of DbpA and B has revealed a critical role in vascular transmigration and as an endothelial potentiator (Tan et al., 2023). Our observation that DbpA (but not DbpB) displays binding to activated endothelium represents an interesting expansion of its functional role and separates functionality of the two members of this paralogous family.

### Description of adhesins binding to pre-activated endothelium

**BBA04**

BBA04, also known as S2, is a surface-exposed lipoprotein (Feng et al., 1995; Barbour et al., 2008). More recently, BBA04 has been identified among cell envelope–associated genes regulated by the second messenger c-di-GMP, linking its expression to environmental sensing and surface remodeling during enzootic transitions (Caimano et al., 2015). In addition, *bba04* is upregulated following tick passage, indicating a potential role during vector-to-host transmission (Adusumilli et al., 2010). The above characteristics suggest that BBA04 may contribute to modulation of the spirochete cell envelope in ways that support host–pathogen interactions during early infection. Here we have provided direct evidence for BBA04 as shear flow-resistant adhesin.

**BBK53**

BBK53 is a surface-exposed lipoprotein that belongs to paralogous family 52 (PF52) and is expressed as part of the RpoS-regulated mammalian-phase transcriptional program (Caimano et al., 2007; Iyer et al., 2015). Deletion of *bbk53*, either alone or together with related PF52 paralogs, does not measurably impair murine infectivity or completion of the enzootic cycle, a phenotype possibly related to extensive functional redundancy within the paralogous family rather than dispensability of the protein (Groshong et al., 2024). To date, no direct host ligand or adhesion target has been identified for BBK53, however its decreased binding to activated endothelium suggests that it may be a GAG binder (**Fig. 5C**). Our experimental approach - expressing BBK53 in a non-adherent heterologous strain lacking confounding paralogs – has allowed functional isolation of BBK53 activity and overcomes a central limitation of prior genetic studies. Using this reductionist system, we demonstrate that BBK53 is sufficient to confer an adhesive phenotype that is shear force-resistant, providing the first direct functional evidence supporting its classification as an adhesin. Unexpectedly, activation of the endothelium results in a decrease in interactions with BBK53 (**Fig. 5C**), a surprising example of the complexity of the adhesion of *B. burgdorferi* to the host vasculature. This may result from a loss of the ligand(s) upon endothelial activation or the presence of new endothelial surface proteins that obscure access to the ligand(s).

**BBK07**

BBK07 is a lipoprotein of *B. burgdorferi* encoded on the linear plasmid lp36 that has been identified as part of the RpoS-dependent regulon activated during mammalian infection

(Caimano et al., 2007). Transcriptomic analyses indicate that *bbk07* expression is low under standard in vitro conditions but is upregulated under conditions associated with mammalian host adaptation (Coleman and Pal, 2009). BBK07 has been shown to be immunogenic during natural infection, eliciting strong antibody responses in infected humans and animal hosts (Barbour et al., 2008; Coleman and Pal, 2009) . Subsequent studies mapped immunodominant epitopes within BBK07 and evaluated its potential utility in serodiagnostic assays for Lyme disease (Coleman et al., 2011). Genetic variation in the presence and sequence of *bbk07* amongstrains has also been reported and may influence host antibody responses (Baum et al., 2012). Despite these findings, no defined host ligand has been identified for BBK07, and its functional role during transmission or dissemination has not been experimentally demonstrated. Here we classify BBK07 as a shear force-resistant adhesin, binding to the endothelium before activation.

**BBA65**

BBA65 is part of a wider family of surface proteins that enhance virulence of a weakly pathogenic spirochete allowing the bacterium to efficiently colonize various tissues during its lifecycle (Adusumilli et al., 2010). The *bba65* gene is upregulated by RpoS during the spirochete's passage through its tick vector, suggesting that the protein may enhance virulence following transmission to mammalian hosts (Caimano et al., 2007; Adusumilli et al., 2010), however its deletion along with *bba64-bba66* does not alter mouse infectivity (Maruskova and Seshu, 2008). Here we characterize BBA65 as a shear force-resistant adhesin, binding to the endothelium before activation.

**BBK32**

Among the best-characterized *B. burgdorferi* adhesins is the fibronectin-binding protein BBK32. We have previously observed transient interactions in real-time between *B. burgdorferi* under shear force in post-capillary mouse venules mediated by BBK32 (Norman et al., 2008; Moriarty et al., 2012) BBK32 binds to both fibronectin and GAGs (Probert and Johnson, 1998; Probert et al., 2001; Fischer et al., 2006) through distinct binding domains (Moriarty et al., 2012). Fibronectin binding occurs through a catch-bond mechanism (Ebady et al., 2016; Niddam et al., 2017) and can induce conformational changes in fibronectin (Harris et al., 2014; Liang et al., 2016). It also binds to the C1r protease subcomponent of the complement system (Garcia et al., 2016), confers bloodstream survival in mice (Caine and Coburn, 2015) and results in attenuated infectivity when absent (Seshu et al., 2006; Hyde et al., 2011). Our recovery of BBK32 (which we previously characterized as a shear force-resistant adhesin (Norman et al., 2008; Moriarty et al., 2012)) in the functional gain screen reported here demonstrates the validity of the methodology used here to identify new adhesins.

**BB0844**

BB0844 is a chromosomally encoded protein in *B. burgdorferi* B31but is usually found on linear plasmids (Banik et al., 2011). Its chromosomal location is likely the result of a stabilized ResT-mediated fusion event (Chaconas, 2005; Kobryn and Chaconas, 2005). Expression of the protein is RpoS-regulated, and it is not required for pathogenicity or maintenance in the tick (Banik et al., 2011). BB0844 is a member of Paralogous Family 12, a family of five DNA-binding proteins (Brangulis et al., 2024). The protein appears to be localized to the periplasm, an unexpected location for a DNA binding protein and has sequence similarity to the coiled coil domain of SMC (structural maintenance of chromosomes) family members (Brangulis et al., 2024)

**ErpK (BBM38)**

ErpK is a surface lipoprotein and a member of the OspF protein family found on cp32 plasmids and is encoded on cp32-6 (Stevenson and Brissette, 2023). Members of the OspF protein family are synthesized during mammalian infection and some members of the family have been shown to bind heparan sulfate, a GAG abundant on cell surfaces and within the extracellular matrix (Brissette et al., 2008). Some have also been selected as adhesins by phage display experiments (Antonara et al., 2007). The entire set of Erps appear to be expendable for murine infection, as a *B. burgdorferi* strain lacking its entire set of cp32 plasmids retains full infectivity (Hillman et al., 2025) and at present their function remains unknown. Here we present evidence for shear force-resistant binding of ErpK to microvascular endothelium.
Moreover, like BBK53 and OspC, we show that endothelial activation reduces the level of endothelial interaction (**Fig. 5C**).

**OspC**

OspC synthesis is induced in the tick following a blood meal (Schwan et al., 1995; Schwan and Piesman, 2000). The protein is an essential early infection factor expressed during initial mammalian colonization and is required for establishment of infection (Grimm et al., 2004; Stewart et al., 2006; Tilly et al., 2006). OspC exhibits antiphagocytic activity (Carrasco et al., 2015) and promotes bloodstream survival by binding to the complement factor C4b (Caine and Coburn, 2015; Caine et al., 2017). The protein has also been reported to bind to plasminogen

(Lagal et al., 2006), fibrinogen (Bierwagen et al., 2019), fibronectin, and or the GAG dermatan sulfate (depending upon OspC) source (Lin et al., 2020). OspC interacts with vascular endothelium (Antonara et al., 2007) and plays a role in tissue tropism and joint invasion (Lin et al., 2020; Tan et al., 2023). Here, we present evidence of direct shear force-resistant interaction of OspC in post-capillary venules prior to endothelial activation. Moreover, we show that endothelial activation reduces the levels of interaction with OspC (**Fig. 5C**) as noted above for BBK53.

**Table S1: *B. burgdorferi* strains.**

| Strain | Strain Bkg | Description | Drug | Reference |
|---|---|---|---|---|
| GCB726 | 5A4 NP1 | 5A4 NP1 (*bbe02::kan*) + pTM61*gent,gfp* | Gm$^R$ Km$^R$ | (Moriarty et al., 2008) |
| GCB989 | HB19, high passage | HB19 HP + pTM61*gent,gfp* | Gm$^R$ | J.Coburn Lab |
| GCB2958 | B31-5A4 | High-infectivity, low passage | | (Purser and Norris, 2000) |
| GCB4036 GCB4038 | B31-5A17 | lp25$^-$, lp28$^-$, $\Delta$*bbk32*::strep, $\Delta$*vlsE*$_{A3}$, pTM61*kan,gfp,pncA* | Km$^R$ Sm$^R$ | (Tan et al., 2022) |
| GCB5109 | GCB989 | pTM61*gent,gfp* + pCS1*kan,pncA,rTel, PresT**ospC*** | Gm$^R$ Km$^R$ | This work |
| GCB5114 | GCB989 | pTM61*gent,gfp* + pCS8*kan,pncA,rTel, PresT**dbpB*** | Gm$^R$ Km$^R$ | This work |
| GCB5119 | GCB989 | pTM61*gent,gfp* + pCS5*kan,pncA,rTel, PresT**bba66*** | Gm$^R$ Km$^R$ | This work |
| GCB5124 | GCB 989 | pTM61*gent,gfp* + pCS6*kan,pncA,rTel, PresT**ospF*** | Gm$^R$ Km$^R$ | This work |
| GCB5130 | GCB989 | pTM61*gent,gfp* + pCS17*kan,pncA,rTel, PresT**bb0844*** | Gm$^R$ Km$^R$ | This work |
| GCB5136 | GCB989 | pTM61*gent,gfp* + pCS3*kan,pncA,rTel, PresT**dbpA*** | Gm$^R$ Km$^R$ | This work |
| GCB5141 | GCB989 | pTM61*gent,gfp* + pCS13*kan,pncA,rTel, PresT**bba65*** | Gm$^R$ Km$^R$ | This work |
| GCB5154 | GCB989 | pTM61*gent,gfp* + pCS14*kan,pncA,rTel, PresT**bba07*** | Gm$^R$ Km$^R$ | This work |
| GCB5159 | GCB989 | pTM61*gent,gfp* + pCS15*kan,pncA,rTel, PresT**bba36*** | Gm$^R$ Km$^R$ | This work |
| GCB5165 | GCB989 | pTM61*gent,gfp* + pCS16*kan,pncA,rTel, PresT**bba34*** | Gm$^R$ Km$^R$ | This work |
| GCB5169 | GCB989 | pTM61*gent,gfp* + pCS18*kan,pncA,rTel, PresT**bbk07*** | Gm$^R$ Km$^R$ | This work |
| GCB5171 | GCB989 | pTM61*gent,gfp* + pCS20*kan,pncA,rTel, PresT**bba64*** | Gm$^R$ Km$^R$ | This work |
| GCB5176 | GCB989 | pTM61*gent,gfp* + pCS19*kan,pncA,rTel, PresT**bbk53*** | Gm$^R$ Km$^R$ | This work |
| GCB5183 | GCB989 | pTM61*gent,gfp* + pCS23*kan,pncA,rTel, PresT**bbi42*** | Gm$^R$ Km$^R$ | This work |
| GCB5188 | GCB989 | pTM61*gent,gfp* + pCS22*kan,pncA,rTel, PresT**bbm27*** | Gm$^R$ Km$^R$ | This work |
| GCB5191 | GCB989 | pTM61*gent,gfp* + pCS2*kan,pncA,rTel, PresT**p66*** | Gm$^R$ Km$^R$ | This work |
| GCB5195 | GCB989 | pTM61*gent,gfp* + pCS24*kan,pncA,rTel, PresT**bba04*** | Gm$^R$ Km$^R$ | This work |
| GCB5201 | GCB989 | pTM61*gent,gfp* + pCS33*kan,pncA,rTel, PresT**bbk32*** | Gm$^R$ Km$^R$ | This work |

Antibiotic concentrations used were: Gm$^R$ 100 µg/ml, Km$^R$ 200 µg/ml, Sm$^R$ 50 µg/ml.

**Table S2: Time correlation**

| | NoMCP | | | MCP-1 | | |
|---|---|---|---|---|---|---|
| | r | p | | r | p | |
| 989 | -0.2129 | 0.3674 | ns | -0.5286 | 0.0543 | ns |
| BB0844 | 0.2564 | 0.335 | ns | 0.1387 | 0.5932 | ns |
| BBA04 | 0.07342 | 0.7722 | ns | 0.06925 | 0.7848 | ns |
| BBA07 | 0.2913 | 0.29 | ns | 0.2254 | 0.4162 | ns |
| BBA34 | -0.3412 | 0.196 | ns | -0.03091 | 0.9181 | ns |
| BBA36 | 0.3843 | 0.1573 | ns | -0.09481 | 0.7357 | ns |
| BBA64 | 0.2697 | 0.394 | ns | 0.2001 | 0.3601 | ns |
| BBA65 | -0.2315 | 0.4227 | ns | -0.2735 | 0.3216 | ns |
| BBA66 | -0.5159 | 0.049 | * | -0.1873 | 0.4425 | ns |
| BBI42 | 0.1981 | 0.5134 | ns | -0.3747 | 0.1036 | ns |
| BBK07 | -0.4884 | 0.0784 | ns | 0.1956 | 0.3829 | ns |
| BBK32 | 0.03681 | 0.889 | ns | -0.3906 | 0.1864 | ns |
| BBK53 | -0.04111 | 0.8853 | ns | -0.2496 | 0.4308 | ns |
| BBM27 | 0.165 | 0.5701 | ns | 0.3679 | 0.1605 | ns |
| DbpA | 0.07963 | 0.7535 | ns | -0.3321 | 0.2264 | ns |
| DbpB | 0.3418 | 0.2302 | ns | -0.4107 | 0.0721 | ns |
| OspC | -0.269 | 0.3497 | ns | -0.1884 | 0.4816 | ns |
| ErpK | -0.1252 | 0.6546 | ns | 0.3495 | 0.221 | ns |
| P66 | -0.2254 | 0.4163 | ns | -0.1853 | 0.4616 | ns |

Spearman correlation time vs number or normalized transient interactions.

**Table S3: Primer sequences used for adhesin cloning.**

| Primer name | Sequence (5'- 3') |
|---|---|
| B70 | CATATGAGCCATATTCAACGGGAAACG |
| B71 | AAAGCCGTTTCTGTAATGAAGGAG |
| B3413 | CTTGGTATAATTGAATTAATATGATTAAATGTAATAATAAAACTTTTAAC |
| B3414 | CTCACAATATAAAAAGAGCTTTAGTTATTTTTGCATTTTTCATC |
| B3415 | CTTGGTATAATTGAATTAATATGAAAAAGAATACATTAAGTGCAATATTA |
| B3416 | CTCACAATATAAAAAGAGCTTTAAGGTTTTTTTGGACTTTG |
| B3417 | CTTGGTATAATTGAATTAATATGAAAAGCCATATTTTATATAAATTAATCATATTT |
| B3418 | CTCACAATATAAAAAGAGCTTTAGCTTCCGCTGTA |
| B3427 | CTTGGTATAATTGAATTAATATGAAAATTGGAAAGCTAAATTC |
| B3428 | CTCACAATATAAAAAGAGCTTTATTTCTTTTTTTTGCTTT |
| B3431 | CTTGGTATAATTGAATTAATATGGAGCAACTTATGAATAA |
| B3432 | CTCACAATATAAAAAGAGCTCTATTCTTTTTTATTAGAATCTTT |
| B3433 | CTTGGTATAATTGAATTAATTTGAAAATCAAACCATTAAT |
| B3434 | CTCACAATATAAAAAGAGCTTTACATTATACTAATGTATGCT |
| B3449 | CTTGGTATAATTGAATTAATTTGAATAAAATAAAATTATCAATATTAATAACA |

| | |
|---|---|
| B3450 | CTCACAATATAAAAAGAGCTTTAATTATTATTAAATTTAAATAATGTG |
| B3451 | CTTGGTATAATTGAATTAATGTGTGTGGGAGACGTATG |
| B3452 | CTCACAATATAAAAAGAGCTTTAAAACGAAGCAGATG |
| B3453 | CTTGGTATAATTGAATTAATTTGATGCAAAGGATAAGTAT |
| B3454 | CTCACAATATAAAAAGAGCTTTAAACATTTCCATAATTTTTC |
| B3455 | CTTGGTATAATTGAATTAATATGATAATAAAAAAAAGAGGAC |
| B3456 | CTCACAATATAAAAAGAGCTTTATTCTTCTATAGGTTTTATTT |
| B3466 | CTTGGTATAATTGAATTAATATGAAAAAAAAAAATTTATCAATTTACA |
| B3467 | CTCACAATATAAAAAGAGCTTTATTTACTCGTCTCTAAAA |
| B3468 | CTTGGTATAATTGAATTAATATGAGTAAACTAATATTGGC |
| B3469 | CTCACAATATAAAAAGAGCTTTAATTATTAAAGCACAAATGT |
| B3470 | CTTGGTATAATTGAATTAATATGAGGATTTTGGTTGGCG |
| B3471 | CTCACAATATAAAAAGAGCTTTATGTAGGTAAAATAGAAACT |
| B3472 | CTTGGTATAATTGAATTAATTTGAAGGATAACATTTTGAAAAA |
| B3473 | CTCACAATATAAAAAGAGCTTTACTGAATTGGAGCAAGAA |
| B3474 | CTTGGTATAATTGAATTAATATGAGAAATAAAAACATATTTAAATTATTTTTTG |
| B3475 | CTCACAATATAAAAAGAGCTTTAATTAGTGCCCTCTT |
| B3476 | CTTGGTATAATTGAATTAATATGAGGATTTTGGTTGGCG |
| B3477 | CTCACAATATAAAAAGAGCTTTATGTAGGTAAAATAGGAAC |
| B3478 | CTTGGTATAATTGAATTAATTTGAAAAGAGTCATTGTATC |
| B3479 | CTCACAATATAAAAAGAGCTTTAGTTAATTAACGAATTAAATG |
| B3516 | CTTGGTATAATTGAATTAATATGAAAAAAGTTAAAAGTAAATATTTGGC |
| B3517 | CTCACAATATAAAAAGAGCTTTAGTACCAAACGCCATT |

**Table S4: *E.coli* strains.**

| Strain | Strain Bkg | Description | Drug | Reference |
|---|---|---|---|---|
| GCE4023 | DH5α | pMC171*kan,pncA,rTel* | $Km^R$ | |
| GCE4502 | DH5α | pCS1*kan,pncA,rTel,PresT**ospC*** | $Km^R$ | This work |
| GCE4508 | DH5α | pCS2*kan,pncA,rTel,PresT**p66*** | $Km^R$ | This work |
| GCE4509 | DH5α | pCS3*kan,pncA,rTel,PresT**dbpA*** | $Km^R$ | This work |
| GCE4519 | DH5α | pCS5*kan,pncA,rTel,PresT**bba66*** | $Km^R$ | This work |
| GCE4521 | DH5α | pCS6*kan,pncA,rTel,PresT**erpK (bbm38)*** | $Km^R$ | This work |
| GCE4533 | DH5α | pCS8*kan,pncA,rTel,PresT**dbpB*** | $Km^R$ | This work |
| GCE4553 | DH5α | pCS15*kan,pncA,rTel,PresT**bba36*** | $Km^R$ | This work |
| GCE4559 | DH5α | pCS16*kan,pncA,rTel,PresT**bba34*** | $Km^R$ | This work |
| GCE4561 | DH5α | pCS13*kan,pncA,rTel,PresT**bba65*** | $Km^R$ | This work |
| GCE4566 | DH5α | pCS14*kan,pncA,rTel,PresT**bba07*** | $Km^R$ | This work |
| GCE4570 | DH5α | pCS17*kan,pncA,rTel,PresT**bb0844*** | $Km^R$ | This work |
| GCE4573 | DH5α | pCS18*kan,pncA,rTel,PresT**bbk07*** | $Km^R$ | This work |
| GCE4578 | DH5α | pCS19*kan,pncA,rTel,PresT**bbk53*** | $Km^R$ | This work |
| GCE4481 | DH5α | pCS20*kan,pncA,rTel,PresT**bba64*** | $Km^R$ | This work |
| GCE4592 | DH5α | pCS22*kan,pncA,rTel,PresT**bbm27*** | $Km^R$ | This work |
| GCE4593 | DH5α | pCS24*kan,pncA,rTel,PresT**bba04*** | $Km^R$ | This work |
| GCE4598 | DH5α | pCS23*kan,pncA,rTel,PresT**bbi42*** | $Km^R$ | This work |

Antibiotic concentration used was $Km^R$ 50 µg/ml.

**Video S1 Legend**

Intravital imaging of neutrophil adhesion to the endothelium in BALB/c mice following infection with wild-type GFP-expressing B. burgdorferi (GCB726). Neutrophils are labeled with anti-Ly6G (red) and venules with anti-PECAM-1 (blue). The left panel video shows the uninfected control (PBS-injected mice), the center panel is a representative video at 5 h, and the right panel is a representative video at 20 h post-infection. The videos display a 200-stack acquisition at 15 fps.

**Video S2 Legend**

BALB/c mice were injected with the wild-type strain GCB726 (n = 5), cultured with 1% blood, as a positive control (left panel). For experimental analysis, mice were injected with phosphate-buffered saline (center panel) or MCP-1, 3 h prior to infection with the GFP-expressing VlsE/BBK32 doubly-deficient strain (GCB4036 or GCB4038) cultured with blood (right panel). Vascular interactions were visualized by high–acquisition rate spinning-disk confocal intravital microscopy 5–50 min post-infection. The videos correspond to a 200-stack acquisition at 15 frames per second (fps).

**Video S3 Legend**

The GFP-expressing doubly-deficient strain GCB4036 was incubated in blood as described previously and either directly injected into MCP-1–treated mice (left panel) or preincubated with proteinase K (150 µg/ml) for 1 h prior to infection (right panel). Following infection, transient vascular adhesive interactions were visualized by intravital microscopy using the same acquisition and analysis conditions as in the preceding video. The videos correspond to a 200-stack acquisition rendered at 15 frames per second (fps).
